# Spatio-temporal transformers for decoding neural movement control

**DOI:** 10.1101/2024.04.15.589534

**Authors:** Benedetta Candelori, Giampiero Bardella, Indro Spinelli, Surabhi Ramawat, Pierpaolo Pani, Stefano Ferraina, Simone Scardapane

## Abstract

Deep learning tools applied to high-resolution neurophysiological data have significantly progressed, offering enhanced decoding, real-time processing, and readability for practical applications. However, the design of artificial neural networks to analyze neural activity remains a challenge, requiring a delicate balance between efficiency in low-data regimes and the interpretability of the results. To this end, we introduce a novel specialized transformer architecture to analyze single-neuron spiking activity. We test our model on multi electrodes recordings from the dorsal premotor cortex (PMd) of non-human primates while performing a motor inhibition task. The proposed architecture provides a very early prediction of the correct movement direction - no later than 230ms after the Go signal presentation across animals - and can accurately forecast whether the movement will be generated or withheld before a Stop signal, unattended, is actually presented. We also analyze the internal dynamics of the model by computing the predicted correlations between time steps and between neurons at successive layers of the architecture. We find that their evolution mirrors previous theoretical analyses. Overall, our framework provides a comprehensive use case for the practical implementation of deep learning tools in motor control research.

## 1. Introduction

Recent innovations in brain recording technologies have enabled simultaneous investigation of vast neuronal populations. On one hand, this is leading to the replacement of conventional methods that rely on analyzing the activity of individual neurons with an ensemble approach, providing insights that could not be inferred solely from the modulation of single units [1–19]. On the other hand, the broad spectrum of recording technologies and the resulting abundance of experimental data have posed challenges in developing standardized routines of analysis consistent across species and conditions. In such a plethora of information, machine learning algorithms, especially deep learning ones, are emerging as cutting-edge tools for skimming redundancies and selecting the most relevant variables for complex system’s dynamics [20–25], opening exciting opportunities for analysis and interpretation of such data [26,27]. In particular, their application in cognitive neuroscience has gained significant attention for their appealing flexibility to link neural representations to behavioral outcomes [28].

In motor control research, decoding neural dynamics to predict motor intentions is of long-standing interest [29]. It is widely recognized that motor control is exerted by an intricate network of cortical, subcortical, cerebellar, and spinal structures [14, 30–41], in which crucial regions are the motor and premotor cortices. These structures play a pivotal role in the encoding of neural signals associated with the planning, execution, and inhibition of voluntary movements. It is also established that single neurons in the motor and premotor cortex codify specific parameters of movement intentions such as direction, speed, and type of intended movements [30, 40, 42–49] before their initiation. Machine learning methods have shown potential to achieve remarkable results in decoding behavioral parameters related to movement [2, 4, 17, 50–56]. Beyond the unquestionable scientific knowledge, one of the ultimate goals of these approaches is enhancing technological progress in prosthetic devices for motor-impaired patients. The scope of exploiting neural signals to control robotic devices or computer cursors in brain-machine interface (BCI) systems [45, 57, 58] is aiding individuals in which gestures, speech [13,57,59,60] or even spinal cord transmissions are compromised [61–64]. Crucially, the efficacy of BCIs heavily depends on their ability to decode and predict the user’s intended behavior accurately, as well as on their power to infer the temporal dynamics. Robustly inferring spatial covariance patterns is also critical, as they serve as a proxy for estimating effective connectivity within the measured neural ensembles. Traditionally, the analysis of neural spiking activity with artificial neural networks has involved the use of 1-dimensional convolutional or recurrent networks [21, 65–68]. In the field of deep learning, transformer-based models, which are built on top of the so-called attention layer [69], have quickly replaced the use of convolutions or recurrences in multiple cases, from natural language processing to cellular automata [22] and computer vision [70]. For the reader less familiar with this terminology, we would like to stress that term “attention” is just a name overlay with the neurophysiological mechanism of attention. In the deep learning context, attention maps are visual representations that highlight the parts of the input data that the model deems most important for making predictions. In natural language processing tasks, for instance, attention maps may highlight specific words or phrases in a sentence. In computer vision tasks, they may pinpoint relevant objects or features in an image. In this paper we will use the term “attention” only in the deep learning sense, without implying any direct parallels to neurophysiological attention mechanisms. When looking at neural data, transformers have a number of interesting features. First, different types of data-formats (time series, images) can be analyzed with a unified architecture by proper pre-processing, e.g., via suitable tokenization or embedding stages [71]. Second, transformers are well-suited to forecasting tasks, such as the prediction of behavioral variables, e.g., movement direction or movement generations vs. movement inhibition, by proper masking of the attention layers, an operation that is highly efficient on modern hardware. Third, attention operates through pairwise comparisons of the input elements. The visualization of the resulting *attention maps* inside their layers provides useful information in terms of which parts of the input are interacting across the architecture [72]. Finally, spatio-temporal variants of the basic transformer [73] have popularized the idea of operating through alternating operations on the spatial and temporal dimension of the data via decomposable layers, making the models perform better on spatio-temporal data with a small number of trainable parameters. Although preliminary investigations have shown promise for the analysis of non-invasive EEG data [21], little has been done for the design of efficient, interpretable transformer-based models for analyzing intraparenchimal recordings of neural populations [74]. To this end, in this paper we propose one of the first transformer-based models to analyze single neuron spiking activity in low-data regimes, which is shown schematically in Fig. 1.

**Figure 1.**
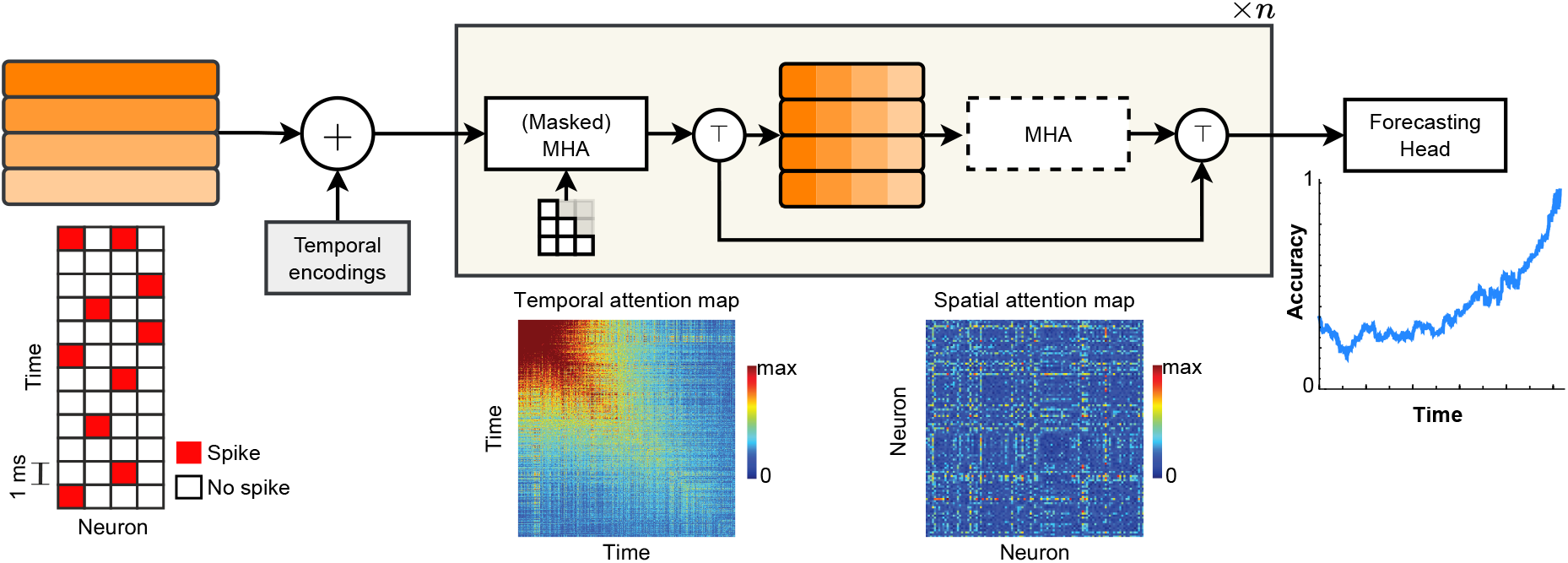
Overall schema of our framework. The experimental single-trial binary data matrix, with dimensions (neurons x time), is pre-processed into a set of vectors (tokens), each representing a small time interval. Having two layers, one temporal and one spatial, a token represent either “one neuron in time” or “all neurons at a given instant” depending on the layer. After adding positional embeddings, we process the tokens via *n* trainable transformer blocks. Each block is composed of multi-head attention (MHA) over the tokens, optionally masked for causal models, followed by another optional block that performs MHA over the transposed tokens. On the bottom of the figure, we show some examples of the outputs we can obtain by visualizing the internal states of the model at each layer. See 2.5 for details.

## 2. Methods

### 2.1 Subjects

Two male rhesus macaque monkeys (Macaca mulatta, Monkeys P and C), weighing 9 and 9.5 kg, respectively, were employed for a Stop signal task, as shown in Fig. 2. Animal care, housing, surgical procedures and experiments conformed to European (Directive 86/609/ECC and 2010/63/UE) and Italian (D.L. 116/92 and D.L. 26/2014) laws and were approved by the Italian Ministry of Health. Monkeys were pair-housed with cage enrichment. They were fed daily with standard primate chow that was supplemented with nuts and fresh fruits if necessary. During recording days, the monkeys received their daily water supply during the experiments.

**Figure 2.**
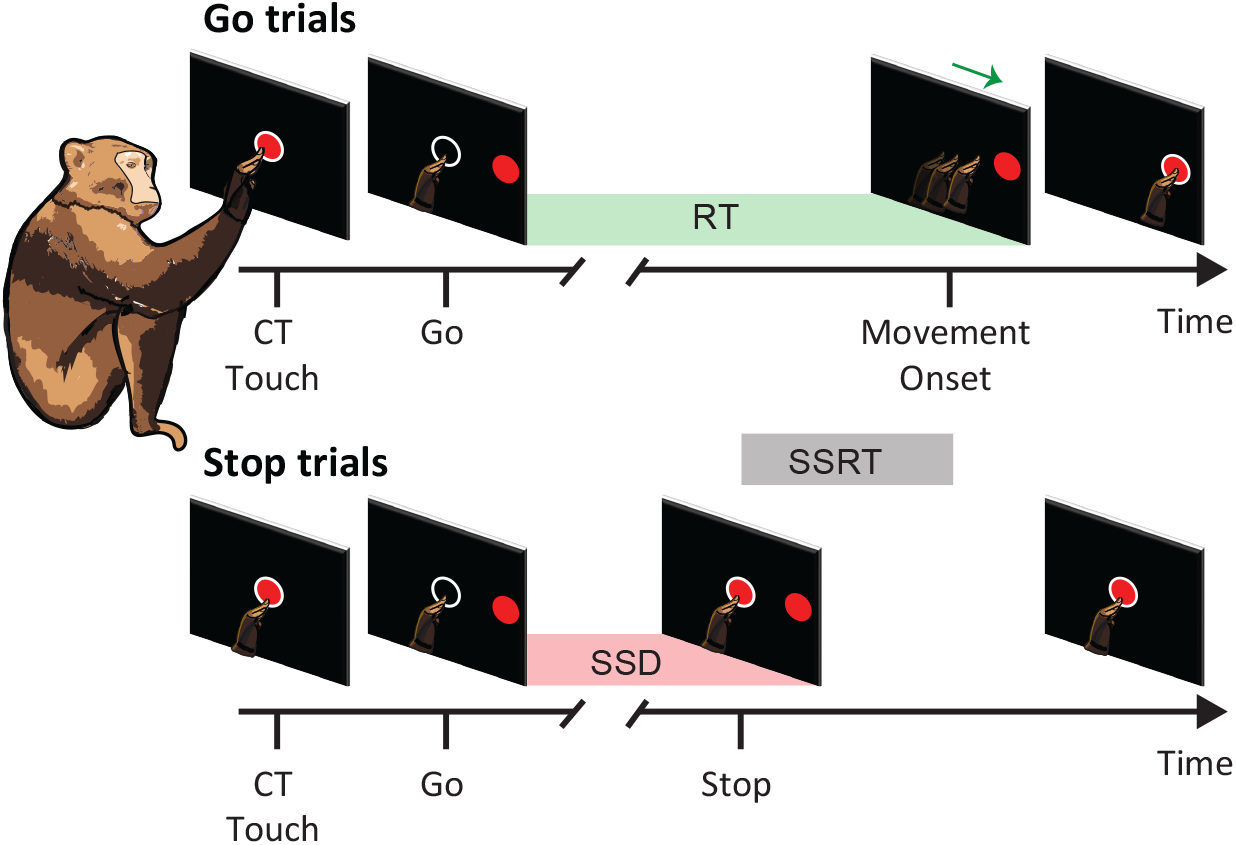
Stop signal task. The target could appear in one of two directions (left or right, D1 or D2). In Go trials, monkeys had to reach and hold the peripheral target to get the reward. In Stop trials, monkeys had to refrain from moving to obtain rewards. Go and Stop trials were randomly intermixed during each session. See Section 2 for further details. RT: reaction time; CT: central target; Go: go signal appearance; M_on: movement onset. SSD, stop signal delay; SSRT, stop signal reaction time. White circles are feedbacks for the touch provided to the animals.

### 2.2 Apparatus and task

The monkeys were seated in front of a black isoluminant background (*<* 0.1*cd/m*2) of a 17-inch touchscreen monitor (LCD, 800 × 600 resolution) inside a darkened, acoustic-insulated room. A non-commercial software package, CORTEX (http://www.nimh.gov.it), was used to control the presentation of the stimuli and the behavioural responses. Fig. 2 shows the scheme of the task. The appearance of a central target (**CT**) (red circle with a diameter of 1.9 cm) signalled the start of the trial. Then, after a holding time of variable duration (400–900 ms, 100 ms increments), a peripheral target (**PT**) (red circle, diameter 1.9 cm) randomly appeared in one of two possible locations on the horizontal plane with respect to the CT (left/right, D1/D2) while the CT disappeared (Go signal). Following the Go signal, the subjects had to reach and hold the PT for a variable time (400–800 ms, 100 ms increments) to receive a reward. The time between the presentation of the Go signal and the onset of the hand movement (**M_on**) is defined as reaction time (**RT**). Stop trials were identical to the Go trials until the Go signal, and then, after a variable delay known as the Stop signal delay (**SSD**), the CT reappeared, signaling the Stop signal. The monkeys had to hold the CT until the end of the trial, which was between 800-1000 ms, to get rewarded for a correct Stop trial (**CS**). To determine the duration of the SSDs a staircase tracking approach was utilized to obtain a success rate of roughly 50% in Stop trials. The SSD rose by 100 ms in the following Stop trial if the monkey successfully inhibited the movement. It was instead reduced by 100 ms if the monkey failed to cancel the response. A Stop trial was deemed incorrect (wrong Stop trial, **WS**) if the monkey removed its hand after the Stop signal and before the trial ended, and no juice was provided. For both correct Stop and correct Go trials, the same amount of reward was given. Stop trials accounted for approximately 25% of all Monkey P trials and 32% of Monkey C trials. White circles after touching the screen were provided to the animals as feedbacks.

### 2.3 Behavioral analysis

In the stop-signal task, the efficiency in movement suppression can be calculated through the so-called Stop Signal Reaction Time (**SSRT**). The expected behavior in the Stop trials is described by the *race model* [75], which comprises the competition towards a threshold of two stochastic processes: the GO process, triggered by the Go signal and whose duration is represented by the RT, and the STOP process, whose duration must be computed. When the GO process wins the race versus the STOP process, movement is generated, resulting in wrong Stop trials. Conversely, when the STOP process prevails, the movement is inhibited, resulting in correct Stop trials. SSRT may be estimated by exploiting the race model by considering the duration of the GO process, the probability of response, and the SSDs). Among various methods to estimate the SSRT, the so-called integration approach has been demonstrated to be the most reliable one [76]. This method considers the existence of a certain probability of response (denoted as P(response)) for a given SSD in stop trials. The longer the SSD, the higher the P(response), indicating a higher probability of a wrong Stop trial. By integrating P(response) over the distribution of Go trials, the nth value Go RT is derived. Subtraction of the SSD from the nth Go RT yields the SSRT. To visualize the procedure, imagine an SSD where P(response) is exactly 0.5. For such an SSD, the Go and Stop processes would have the same probability of winning the race, meaning that they would both reach the threshold at the same time with equal probability. The ending times of the Stop and Go processes are the end of the SSRT and the end of the RT respectively, i.e., the beginning of the movement. Behavioural results for the analyzed sessions (2, one for each animal) are reported in Table 2.

### 2.4 Extraction and processing of neuronal data

A 96-electrode array (Blackrock Microsystems, Salt Lake City; electrode spacing: 0.4 mm) was surgically implanted in the left dorsal premotor cortex (PMd) taking as reference the arcuate sulcus and pre-central dimple. The raw signal sampled at 24.4 kHz (Tucker Davis Technologies, Alachua, FL) was acquired. We used the spike sorting toolbox KiloSort3 with the following settings to get single neuron activities from the raw signal, similar to previous work [14, 77]. Thresholds for template-matching on spike detection: (9 9); Lambda: 10 (bias factor of the individual spike amplitude towards the cluster mean); Area Under the Curve split: 0.9 (threshold for cluster splitting); Number of blocks: 5 (the number of blocks channels are divided into for estimating probe drift). The output of the sorter was curated manually in Phy (v2.0; 17) to merge clusters erroneously split by the automated sorter. Then, a binary data matrix of dimensions (neuron x time) with a time resolution of 1 ms (1 for a spike, 0 for no spikes) was obtained for each single trial of each monkey.

### 2.5 Transformer architecture

We use a customized transformer architecture [69] arranged in a space-time configuration [73] that is tailored to simultaneously predicting movement direction (D1 vs. D2), movement intentions (movement vs. inhibition), and can estimate the spatial and temporal correlations between the recorded neurons, using as input only their experimental spiking activity. Fig. 1 provides a visual schema of our model, denoted spatio-temporal transformer (**ST-T**), and its outputs. First, a single-trial binary spike data matrix is preprocessed to convert it to a sequence of fixed-dimensional vectors (which we refer to as *tokens*). Then, we process the sequence via alternating attention-based blocks that operate on the time or on the neuron dimension of the input data. The network is trained for each time step to predict a predefined behavioural event, e.g. the movement onset. We show that this design provides at the same time a strong predictive accuracy of the event, as long as a set of explainable maps that describe the dynamics of the interactions between time steps and neurons inside each intermediate layer of the model, which can be visualized and compared to experimental ones and/or to theoretical models to validate the performance. By looking at the accuracy of the predictions and the evolution of the internal states of the network, several insights can be gathered, which are summarized qualitatively here and quantitatively in the experimental section.

#### 1) Nonlinear correlation analysis

It has been shown that temporal and spatial correlations among neurons carry non-trivial features of the dynamics of executed and inhibited movements [14, 19, 41, 77–82]. Even if pairwise correlation can be computed with standard statistical techniques [83], it is not guaranteed that they are indicative in terms of predictive power for movement parameters, i.e., the type and direction. Because (artificial) neural networks are instead trained to evolve their internal representations to maximize performance, looking at the correlations across the internal embeddings of neurons and time intervals, information is extracted in the form of nonlinear, task-dependent correlations. We show below that suitably designed transformer models are a natural fit for this assignment, since representations are updated by pairwise comparisons of the internal tokens. The resulting *attention maps* can be immediately visualized to understand the evolution of these dynamics (see Section 2.5). In addition, the same model can be customized to work across different axes (time and/or space), providing both attention maps as auxiliary outputs (see the bottom plots in Fig. 1 for examples).

#### 2) Forecasting accuracy

Second, operations inside a transformer can be easily masked to obtain *causal* variants, in which the output *y*_*t*_ at time *t* does not depend on input data **x**_*>t*_ corresponding to subsequent time indexes. By training a separate predictor on each output *y*_1_, …, *y*_*T*_ and looking at the average accuracy of the predictions, we can analyze when, after the presentation of the Go signal, we can accurately predict the trial outcome and the direction of movement, independently of the specific RT and SSD. We now briefly describe the models that we used in the analysis. For the D1 vs. D2 prediction, data is extracted for each trial in a fixed interval of 400 ms, i.e., from 100 to 500 ms after the Go signal presentation, resulting in a set {**x**_*t*_}, **x**_*t*_ ∈ ℝ ^*N*^ corresponding to *N* neurons and *t* = 1, …, *T* time instants (*T* = 400). For the movement vs. inhibition prediction, data is extracted for each trial from 100 to 300 ms after the Go signal presentation, to control for the stop Signal appearance (T=200ms) (see Section 3). Because transformer models are equivariant to permutations, sinuisodal embeddings [69] are added to the initial embeddings to encode information about the distance between temporal instants. As shown in Fig. 1, embeddings are then processed by a series of *n* trainable blocks. Each of these blocks contains up to 2 standard transformer blocks which process data across the two axes. First, a temporal transformer block is applied:

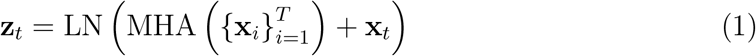

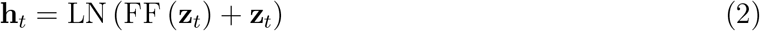

where LN stands for layer normalization, SA for self-attention, and FF is a two-layer fully-connected network applied to each token independently [69]. Note that the only operation which combines information from multiple tokens is the SA. Denoting generically by {**x**_*i*_} the set of input tokens, MHA proceeds by first linearly projecting each of them three times with different trainable matrices to obtain query tokens **q**_*i*_, value tokens **v**_*i*_, and key tokens **k**_*i*_, before combining them as:

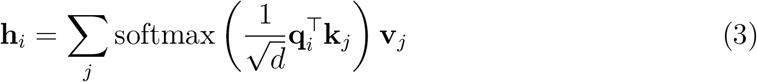

The outputs of the softmax operation can be combined into a *T × T* matrix, that we call the *temporal attention matrix*. We also consider a causal variant of this layer, in which all values such that *j > i* are masked by setting the corresponding dot product to −∞. In this way, each output token **h**_*t*_ will only depends on the input tokens 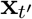 such that *t*^*′*^ ≤ *t*, preserving the temporal dynamics of the signal. For the non-causal variant, we also include a second, symmetrical block working on the channel-wise (spatial) axis. Denoting by **H** ∈ ℝ^*T ×N*^ the matrix collecting the output of the first block, we apply a second block by first transposing the matrix, which is equivalent to considering the columns of **H** as the input tokens, before transposing again for the next block. We call the corresponding attention map of size *N ×N* the *spatial attention matrix*. Note that SA can be extended by considering a multi-head variant (with multiple SA blocks running in parallel), but we ignore it here for better interpretability of the resulting attention maps. For training, we apply a prediction head to each output token independently, obtaining a series of predictions *ŷ*_1_, …, *ŷ*_*T*_. For the non-causal variant, we discard all predictions except the last one, which we train to predict the binary label corresponding to the chosen event (D1 vs. D2 or movement vs. inhibition). For the causal variant (Causal ST-T), we consider a separate classification task for each time interval and we optimize by minimizing the average loss across intervals. The test accuracies *a*_*t*_ for each interval can be plotted to visualize the moment in which the behavior becomes successfully predictable by the model.

### 2.6 Comparison with other models

We benchmarked the ST-T’s performance against various alternatives: a CNN architecture, a logistic regression model (LR), a support vector machine with a Gaussian kernel (G-SVC) and a correlation coefficient-based decoder (CC-D). Both the LR and the G-SVC are trained with the scikit-learn [84] implementation using a grid-based search for optimizing the hyperparameters using the same validation and test sets of the CNN and the ST-T. The hyperparameters of the ST-T and CNN architectures are reported in Table 1. For the CC-D we used some customized functions of the NDT Matlab toolbox [85].The classifier was trained to discriminate among movement direction or movement generations vs. movement inhibition. The classifier employed a cross validation approach, which required splitting the trials into *k* splits and using *k* – 1 data splits for training the classifier while using the last remaining split for testing. This procedure was repeated k times using a different test split every time. Finally, all these steps were repeated 50 times while resampling the training and the test splits within the dataset. The prediction accuracy of the classifier is reported as the average classification accuracy across all the resampling runs. The value of k was chosen as a number less than or equal to the minimum available trials across conditions.For the sake of a direct comparison with the ST-T classification accuracies and to assess the overall decoding performance, we performed the population decoding analysis using non-overlapping bins 1 ms and 10 ms respectively.Tables with the results of comparisons between all models across conditions and monkeys can be found in the Supplementary Information (SI).

**Table 1.** Setting of hyperparameters for different predictions.

| Hyperparameters | D1 vs. D2 |  |  | Movement vs. Inhibition |  |
| --- | --- | --- | --- | --- | --- |
|  | CNN | ST-T | Causal ST-T | CNN | ST-T |
| Heads | - | 1 | 1 | - | 1 |
| Layers | - | 5 | 2 | - | 2 |
| Learning Rate | $1e^{-4}$ | $1e^{-4}$ | $1e^{-4}$ | $1e^{-4}$ | $1e^{-5}$ |
| Epochs | 15 | 80 | 150 | 30 | 80 |
| Batch size | 8 | 8 | 8 | 8 | 8 |

**Table 2.** Behavioral results (mean ± std) for the analysed recording sessions.

| Subject | RT | SSD | SSRT |
| --- | --- | --- | --- |
| Monkey P | 595 ± 90 ms | 343 ± 70 ms | 209 ms |
| Monkey C | 671 ± 111 ms | 423 ± 128 ms | 207 ms |

## 3. Experimental results

We performed single-trial analyses aligning the time series of neural activity to the Go signal for each session of both monkeys. Such alignment was motivated by the evidence that, after the Go signal, PMd neurons express strong modulations associated with movement control. Indeed, it has been shown that the motor planning of actions in the PMd is encoded at the population level through synchronization patterns that significantly peak around 200 ms before movement initiation [14,19,30,34,40,41,78,79,81, 82,86,86–90]. Here our goal is to propose an effective method to decode motor intentions from neural activity before the actual movement is either executed or inhibited, i.e. in the preparatory phase (before M_on). In the jargon of BCIs this would translate in establishing a baseline open-loop performance in an off-line setting. Therefore, considering only the motor planning phase, our analysis cannot be influenced by visual feedback adaptation of the movement trajectory, which is known to be a crucial factor requiring careful calibration in closed-loop settings [56, 58]. Moreover, we also do not analyze/include somatic/visual feedback signals, which in closed-loop paradigms are usually provided to the subject to tune and refine the control of external devices (e.g., robotic arms, computer cursors etc..). It is also true that visual stimuli can modulate neural activity in motor and premotor areas [91–95]. However, only a low percentage of neurons in the motor cortices encode purely visual information, with such an effect resolving around 55–70 ms after visual stimulation [92, 94–100]. In our case, this would correspond to the visual encoding of the target position. To eliminate the influence of this early visual information on the performance, we excluded from our analysis the first 100 ms after the Go signal. Hence, we are confident that our results account for only the motor planning-related activity of premotor neurons, well in advance of the time when the arm, moving, could provide somatosensory and visual feedbacks to the system. Considering the RT distributions shown in Table 2, for the D1 vs. D2 prediction, we aligned neural activity [+100, +500] ms to the Go signal for both monkeys. Tables with condition comparisons between the ST-T and the tested alternatives can be found in the Supplementary Information (SI). 10

### 3.1 Prediction of movement direction

The accuracy of the predictions for the ST-T is reported in Figure 3 for both monkeys, and trial types (Go and correct Stop trials). For all the other methods see Tables S1, S4, S5. Overall, ST-T performs better than the other models, with an accuracy of over 90% in almost all sessions. The CNN and the others methods are less consistent, yielding a high accuracy for Monkey P (*>* 94%) but not for Monkey C (≤ 70%). This was somewhat expected, given that our ST-T did not perform (relatively) well for this prediction and given the stereotypical and low-noise neural dynamics of Monkey P [14, 77], which enables even simpler and linear non-causal models to achieve efficient classification. The good performance of LR and G-SVC can also be understood by considering some previous results of our group, in particular Pani et al. 2022 [14]. In this work, we showed that we could project high-dimensional neural activity in PMd during movement planning and inhibition into a low-dimensional state space, identifying two key axes: the Holding-and-Planning Axis (HPA) and the Planning-and-Execution Axis (PEA). HPA refers to slow ramp-like changes in neural activity associated with motor planning, whereas PEA refers to rapid, threshold-like changes linked to movement execution. The HPA corresponds to a “holding plane” that neural activity occupies before movement generation, within which activities are confined when a movement is inhibited. Activity escapes this plane into a distinct, orthogonal subspace, demonstrating a clear separation between planning and execution at the population level and initiating movements. During the planning phase, the neuronal dynamics along the HPA exhibit a relatively linear ramp-up of activity, suggesting that the build-up of motor planning signals is linear over time. The transition from the HPA to PEA marks the transition from planning to execution, with the dynamics along the PEA involving a threshold-like mechanism and the movement execution phase corresponding to a quick crossing of this threshold.

**Figure 3.**
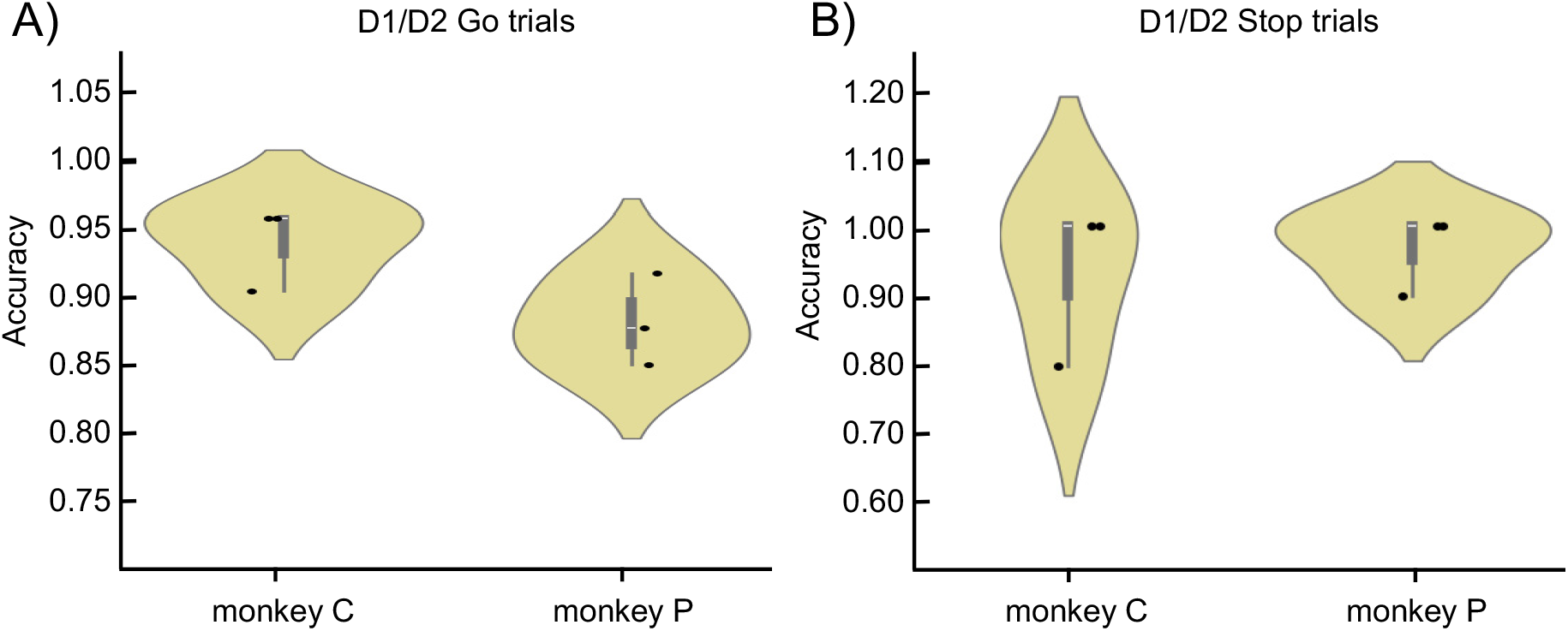
Accuracy distributions over model runs for D1 vs. D2 prediction for Go (panel A) and correct Stop trials (panel B) using the ST-T for both monkeys . For comparison with the other models see Table S1 and S3.

It is worth noting that our model predicts in advance the correct direction of movement for correct Stop trials as well, where movement is eventually inhibited. This is relevant for two reasons. The first is that it strongly confirms, at the neural level, the assumptions underlying the Stop Signal paradigm and its effectiveness in the study of response inhibition: that the movement is planned and then inhibited also in the correct stop trials (see also Section 2.3). The second is that, our model provides an effective method to quantitatively measure how motor planning fully develops even when the movement is not eventually generated. These are two basic premises for being able to address the movement vs. inhibition prediction (see next paragraph).

We tested the stability of our prediction exploiting the Causal ST-T. In this way, we could obtain the time evolution of the prediction accuracy for movement direction in Go trials, as shown in Figure 4. In Figure 4 we also compared the causal ST-T accuracy to the CC-D’s accuracy over time. When considering the temporal coding properties of the population activity, the CC-D’s performance was significantly worse failing to achieve comparable time resolutions and accuracy. Our results are especially significant because the CC-D is one of the most employed methods in neurophysiology studies for the estimation of temporal population coding related to cognitive and motor processes [51,85,101–108]. In this regard, we would like to emphasize that CC-D achieves above-chance performance (but still ≤ 67%) only by significantly increasing the time bin of analysis (see Tables S5 and S6). This, while demonstrating the method’s effectiveness, also highlights its inherent limitations in describing fine time scales, thereby significantly restricting its application domain.

**Figure 4.**
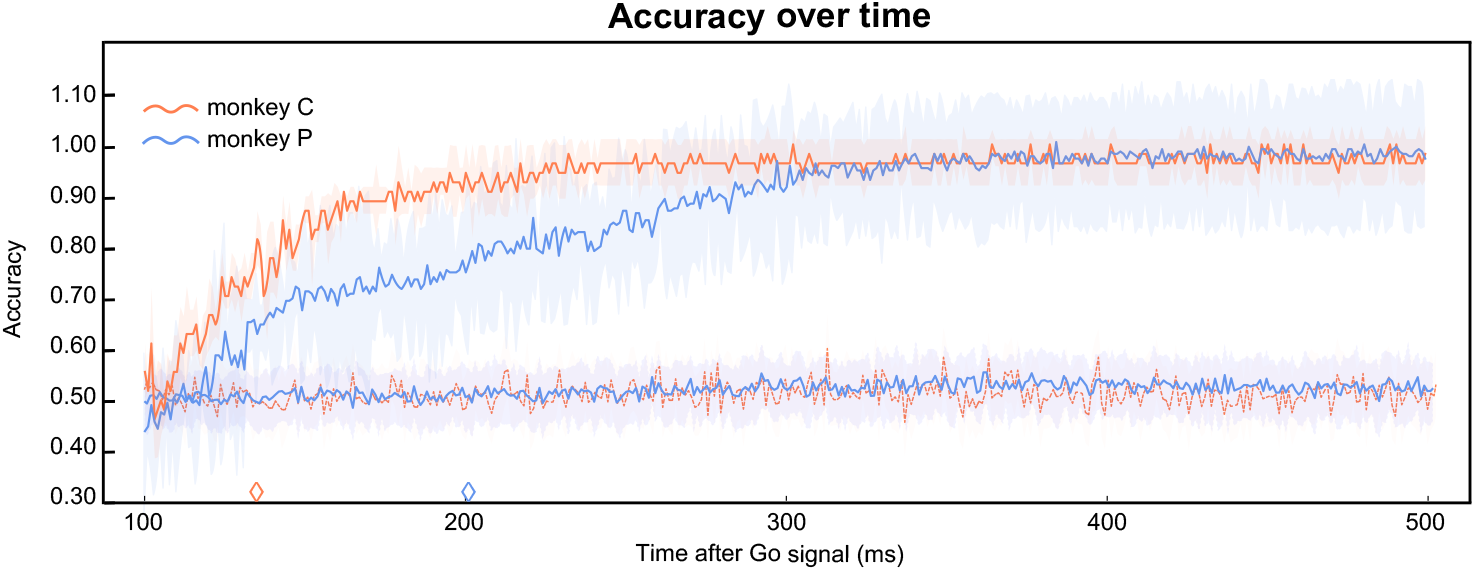
Time evolution of the accuracy for D1 vs. D2 prediction for all the Go trials compared between the Causal ST-T (solid lines) and the CC-D (dashed lines) . Shaded areas are the standard deviations over multiple model runs. Diamonds mark the time in which the ST-T reaches an accuracy ≥ 80%. This time point cannot be identified in CC-D, whose performance always remains at chance level for this time resolution.

Using the Causal ST-T, we repeated the analysis first considering all the trials together and successively dividing the trials according to their RTs (four groups, as shown in Figure 5 and Table S2). Considering all the trials together we are able to correctly decode movement direction with an 80% accuracy ∼ 200 ms after the Go signal for monkey P and after ∼ 130 ms for monkey C. The level of accuracy then remains stable after ∼ 230 ms for monkey C and after ∼ 300 ms for monkey P. Since trials with slower RT imply later motor planning, we expect the time at which the prediction reaches high levels of accuracy to be progressively further away from the Go signal as the RT increases. As shown, it is evident how a high accuracy is reached earlier after the Go-signal for trials with faster RTs, confirming our assumptions. This is significant because it demonstrates the ability of our architecture in predicting the behavior efficiently for different levels of motor preparation.

**Figure 5.**
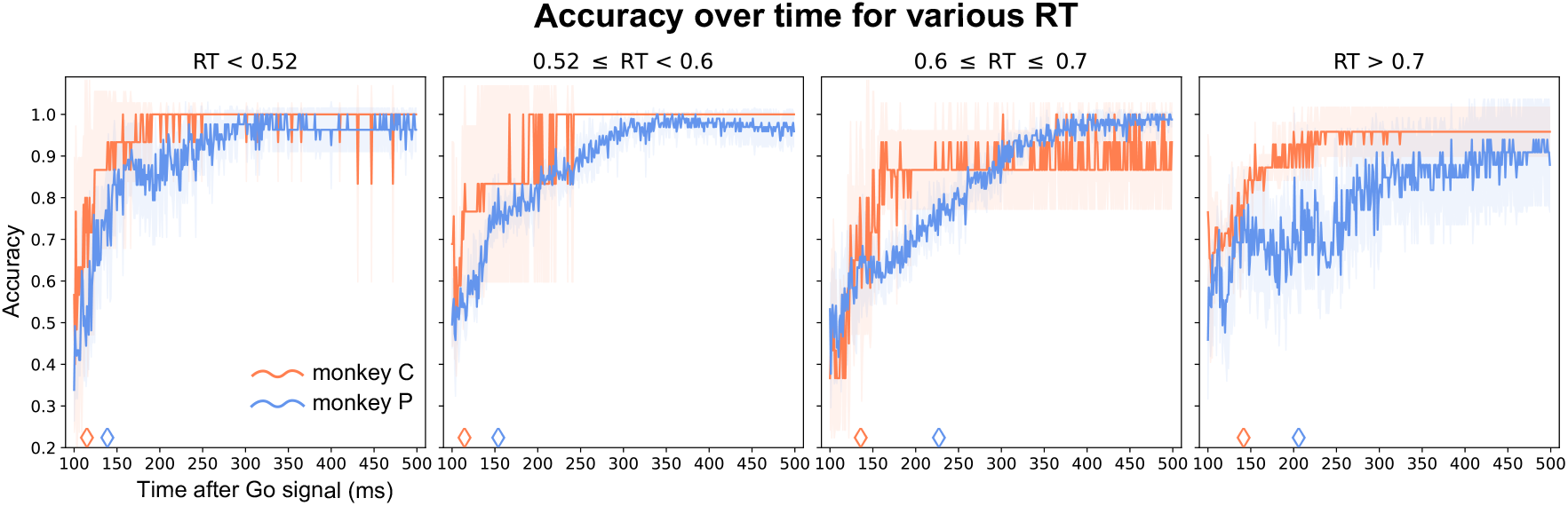
Evolution of the prediction accuracy for D1 vs. D2 prediction for Go trials with different RT for the Causal ST-T. Shaded areas are the standard deviations over multiple model runs. Diamonds mark the time in which accuracy ≥ 80%.

### 3.2 Prediction of movement inhibition

Our goal was also to predict the generation or inhibition of a movement. To do so, we chose the Stop trials corresponding to the SSD with the highest number of Stop trials presented, specifically 300 ms for monkey P and 400 ms for monkey C. This choice was forced by the smaller number of trials available for the correct Stop condition compared to Go trials and led to the inclusion of 62 Stop trials for monkey P and 27 for monkey C. This also prompted us to employ a simpler architecture, with only 2 layers instead of 5, so that the model could attain better generalization without overfitting. For this prediction, we aligned neural activity [+100, +300] ms after the Go signal, thus controlling for the possible bias of including the Stop signal presentation in the time window of analysis. We performed two tests: in the first, we compared only the Stop trials (correct vs wrong) while in the second one we used all three trial types (go, correct Stop and wrong stop trials). Correct Stop trials can be compared with Go trials only if an appropriate subset of the latter is considered. Such subset must include all the Go trials that, if a Stop had appeared, would have had a high probability of being inhibited. These trials are called latency matched trials, and are the trials with *RT > SSD*+*SSRT* (see also Section 2.3). Results are shown in Figure 6. As before, we compared the ST-T performance with that of other methods (see Table S3). Consistent with the D1 vs. D2 prediction, ST-T yields a significantly higher accuracy in both tests. However, the CNN still achieves a very good level of accuracy (an average of 76.8 % across monkeys and tests). We attribute the lower performance for monkey C in the first test to the fewer number of trials available. The results obtained are quite noteworthy, as they demonstrate the efficiency of our model in predicting very early and accurately whether motion will be generated or not, even with very few trials available.

**Figure 6.**
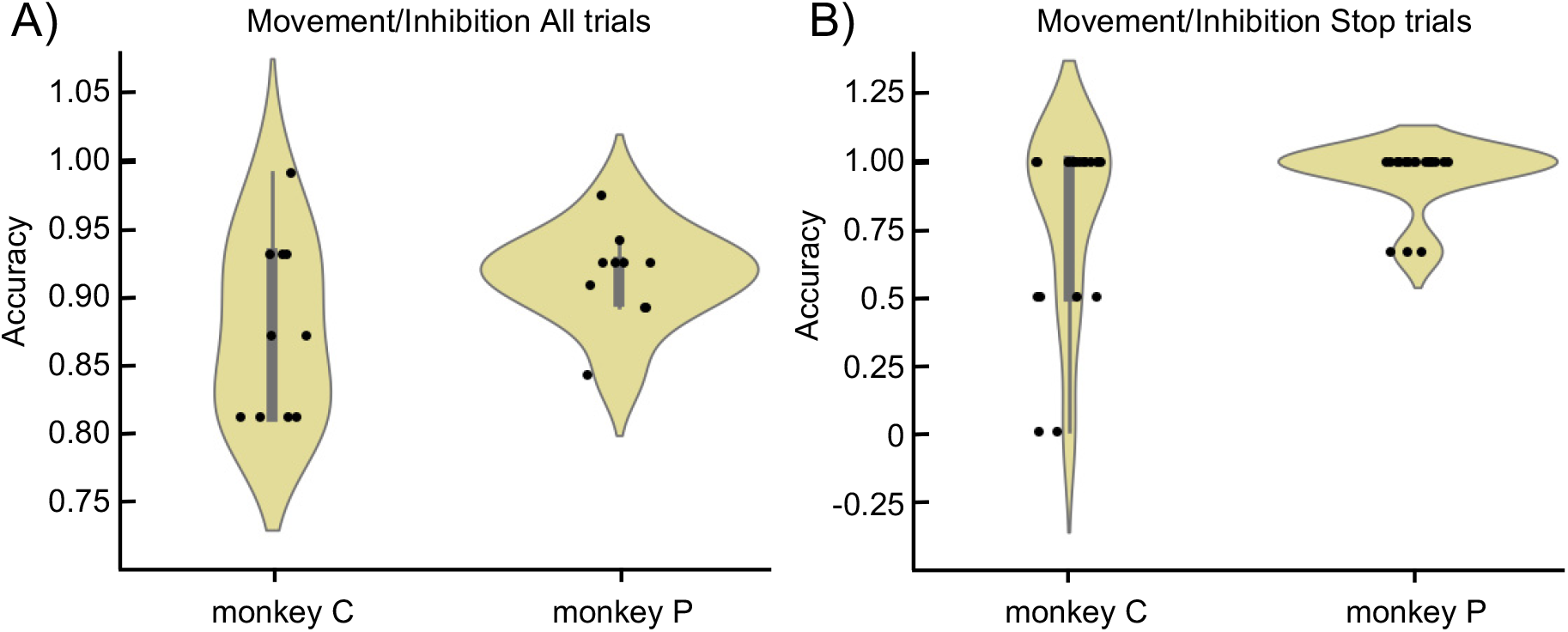
Accuracy distributions over model runs for movement vs. inhibition prediction for all trial types (panel A) and for Stop trials only (panel B) using the ST-T for both monkeys . For comparison with the other models see Table S3.

## 4. Conclusions

In this study we presented a spatio-temporal transformer (ST-T) architecture for decoding neural movement control from multi-electrode recordings *in vivo*. The advantages of the ST-T architecture include its ability to account for possible nonlinear interactions through attention layers, efficiently generalizing with minimal training data, and ensuring causal predictions by maintaining temporal relationships at the millisecond scale. These features, which contrasts with the limitations of the other methods tested, make it very attractive for possible real-time applications. Overall, the ST-T architecture offers better accuracy, interpretability, and efficiency, particularly under conditions of sparse data. One limitation is evident in conditions where neural dynamics are stereotypical and low-noise, such as the D1 vs. D2 prediction for Go trials of monkey P. In this case, simpler models were more accurate, which shows that ST-T doesn’t clearly outperform more straightforward linear models when training data is large, of high quality and may contain linear relationships. Still, fine-tuning the ST-T architecture could improve its performance even in these simpler conditions, but the necessity and advantage are context-dependent. Nonetheless, the efficacy of ST-T emerges when temporal coding is complex or possibly non-linear, as demonstrated by better performance compared to a standard and widespread method such as the CC-D when accounting for fine-grained temporal dynamics. But it is not all about accuracy. The advantages of the ST-T encompasses also its capacity to simultaneously estimate temporal and spatial attention maps. We give here a perspective of how the spatio-temporal attention maps provided by the model can be interpreted in a neurophysiological context. We leave a more extensive formal analysis for future investigations. The temporal attention maps represent how much the temporal correlations among the recorded neurons impact the required prediction. In our case, the magnitude of the matrix elements indicates the importance that the collective state of the neural ensembles at that time instant has in determining the forecast of the behavioral outcome. In Figure 7 panel A we compared the temporal attention matrices for the D1 vs. D2 prediction. The model’s ability to identify new correlation patterns becomes evident as its depth grows, especially in correct Stop trials where condition-specific patterns of synchronization appear already from layer 2. This is particularly noticeable in panel B when looking at the distributions of their matrix elements as a function of depth. Interestingly, until about 250 ms after the Go signal, the synchrony patterns inferred by the model are very similar between conditions. As discussed above, this is suggestive of the partitioning of the phase space of neural activity into common and specific subspaces according to behavioral conditions [14, 78]. This strategy can also be extended to the spatial component of attention maps. Indeed, synchronized firing between spatially separated pairs of neurons is commonly employed as a measure of functional connectivity. As shown in earlier research, differences in the topological organization of spatial synchronizations have a key role in explaining how motor cortices are involved in both movement generation [19, 41, 109, 110] and inhibition [19, 41]. Therefore, spatial attention maps produced by each layer could be compared to the experimental covariance matrices, looking for topological invariants between the two. In this scenario, these matrices could be interpreted as the adjacency matrices of weighted networks, with the spatial attention map representing a naive unsupervised inference of the spatial interaction graph of the system. However, with other depp-learning methods also asymmetric interactions (e.g., directed graph) could be obtained [22, 24, 25]. As also addressed by Ha et al. [22], this hints at how our ST-T architecture can be an alternative route for developing new models for so-called relational-inference problem: detect interactions among components of a dynamical system based on samples of the observed time-series [24, 25, 111–114]. In this regard, very recent works are addressing relational prediction from spike data [23] while others are seeking to infer detailed models and interpretable high-order microscopic theories directly from data [115]. These works exposed how to use deep learning in physical systems where some detail of microscopic interactions is known but effective theories are needed to explain the emergence of multiscale collective behavior. This is perfectly fitting for almost every experimental setting in a behavioral neurophysiology experiment. This not only contributes to making the output of such deep models interpretable, but it also opens up realistic possibilities for using these methods as a powerful unsupervised way of estimating effective connectivity in neural systems.

**Figure 7.**
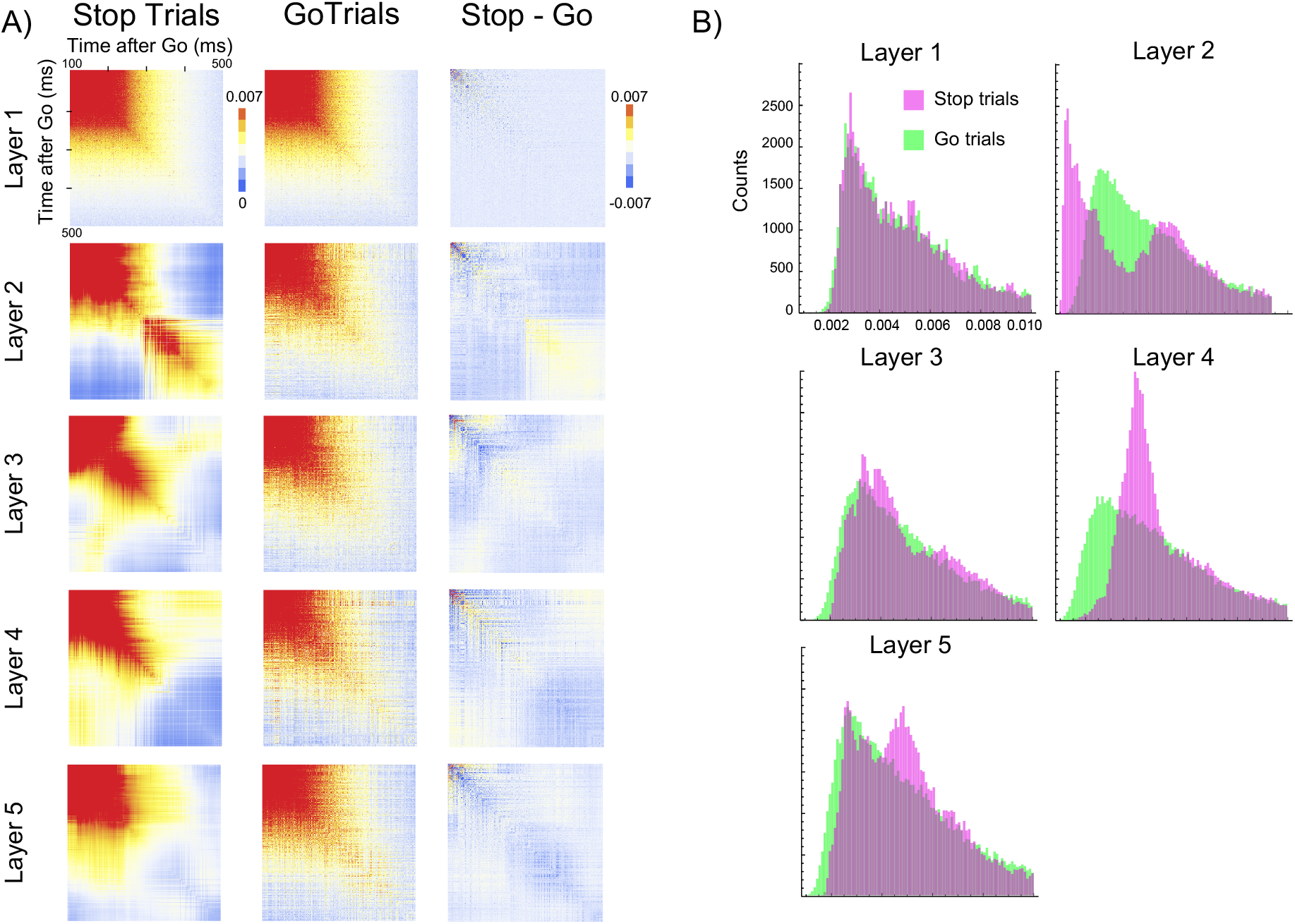
Qualitative comparison between temporal attention maps of Go and Stop trials for the D1 vs. D2 prediction for monkey P. **Panel A)** Attention maps of each layer. Ticks are every 100 ms and mark the time after the Go signal presentation **Panel B)** Distribution of their entries.

Intriguingly, quantitative and formal study of transformers and their attention maps can facilitate the development of a causal connection between findings obtained in diverse disciplines. In this regard, it has recently been demonstrated that the spatial and temporal correlation matrices of a system of neurons, to which maps are strongly related, reflect its potential and kinetic energy contribution, respectively. [77,116] Thus, developing reliable methods for estimating such interaction matrices from empirical recordings can contribute significantly to building coherent physical theories for neural systems. A significant effort in this direction comes from the work of Rende et al. 2024 [117].In this research, the authors demonstrate that a single layer of self-attention in transformers trained via Masked Language Modeling (MLM) learns the conditional probability distributions of a fundamental model of statistical physics, the generalized Potts model. This association strongly supports the interpretation of self-attention training as addressing the inverse Potts problem through the pseudo-likelihood method, a statistical inference approach. Such equivalence provides a clear theoretical framework for understanding the self-attention mechanism in transformers, allowing for the use of well-established statistical physics techniques [117, 118]to analyze and predict transformer behavior.

## Acknowledgments

The project was partly funded by Sapienza grant RM1221816BD028D6 (DESMOS), European Union’s CHIST-ERA programme under grant agreement CHIST-ERA-19-XAI-009 (MUCCA), the PNRR MUR project PE0000013 FAIR, and by a NextGenerationEU grant (ID IR0000011, “EBRAINS-Italy”, Stefano Ferraina).

## Data availability statement

Data are available from the corresponding author(s) upon reasonable request.

