## Supplemental Information for "Spatio-temporal transformers for decoding neural movement control"

**Table 1.** Results for D1 vs. D2 prediction compared between ST-T, CNN, LR and G-SVC. For CC-D results see Tables 4 and 5.

| Subject | CNN | | LR | | G-SVC | | ST-T | |
|---|---|---|---|---|---|---|---|---|
| | Go t. | Stop t. | Go t. | Stop t. | Go t. | Stop t. | Go t. | Stop t. |
| Monkey P | 99.8% | 94% | 96% | 47% | 94.3% | 45.6% | 88% | 96% |
| Monkey C | 70.7% | 66% | 76.5% | 46.6% | 56% | 53.3% | 93% | 93% |

**Table 2.** Results for D1 vs. D2 prediction using Causal ST-T for various gropus of RTs.

| Go trials only | Time after the Go signal to reach >80% of accuracy | | | | |
|---|---|---|---|---|---|
| | Overall | $RT < 520$ | $520 \leq RT \leq 600$ | $600 \leq RT \leq 700$ | $RT \geq 700$ |
| Monkey P | 204 ms | 140 ms | 154 ms | 233 ms | 201 ms |
| Monkey C | 135 ms | 113 ms | 115 ms | 135 ms | 135 ms |

**Table 3.** Movement vs. inhibition prediction results compared between ST-T, CNN, LR and G-SVC. For CC-D results see Tables 6 and 7.

| Subject | CNN | | LR | | G-SVC | | ST-T | |
|---|---|---|---|---|---|---|---|---|
| | All t. | Stop t. | All t. | Stop t. | All t. | Stop t. | All t. | Stop t. |
| Monkey P | 83.9% | 75% | 53.3% | 47% | 60% | 45.6% | 95.1% | 95% |
| Monkey C | 71.4% | 77% | 47.7% | 71.1% | 46.6% | 74.3% | 92.6% | 80% |

**Table 4.** Results for D1 vs. D2 prediction for the CC-D with non-overlapping bins of 1 ms; Monkey P (Go Trials: k=40; Stop Correct trials: k=40), Monkey C (Go Trials: k=40; Stop Correct trials: k=19).

| Subject | Go t. | Stop t. |
|---|---|---|
| Monkey P | 52.13% | 51.39% |
| Monkey C | 51.13% | 51.16% |

**Table 5.** Results for D1 vs. D2 prediction for the CC-D with non-overlapping bins of 10 ms; Monkey P (Go Trials: k=40; Stop Correct trials: k=40), Monkey C (Go Trials: k=40; Stop Correct trials: k=19).

| Subject | Go t. | Stop t. |
|---|---|---|
| Monkey P | 67.17% | 65.4% |
| Monkey C | 65.57% | 66.58% |

**Table 6.** Results for movement vs. inhibition prediction using latency matched trials for the CC-D with non-overlapping bins of 1 ms; Monkey P (k=13), Monkey C (k=5).

| Subject | All t. | Stop t. |
|---|---|---|
| Monkey P | 33.93% | 50.93% |
| Monkey C | 33.51% | 51.69% |

**Table 7.** Results for movement vs. inhibition prediction using latency matched trials for the CC-D with non-overlapping bins of 10 ms; Monkey P (k=13), Monkey C (k=5).

| Subject | All t. | Stop t. |
|---|---|---|
| Monkey P | 36.37% | 53.63% |
| Monkey C | 31.77% | 60.98% |
